# Impact of a native hemiparasite and mowing on performance of a major invasive weed, European blackberry

**DOI:** 10.1101/2022.01.22.477376

**Authors:** Robert M. Cirocco, Jennifer R. Watling, José M. Facelli

**Affiliations:** School of Biological Sciences, The University of Adelaide, Adelaide, SA 5005, Australia; Ecology and Environment Research Centre, Manchester Metropolitan University, Manchester, M15 6BH, UK

**Keywords:** biocontrol, biological agent, chlorophyll fluorescence, parasitic plants, plant invasions, *Rubus*, weed management

## Abstract

1. Plant invasions are a major global threat to biodiversity. Traditional methods of weed control are falling short, and novel and environmentally friendly control tools are needed. Native parasitic plants are showing promise as effective biocontrols for some of the worst weeds, however, their application is in its infancy.
2. First, we established the native parasitic plant, *Cassytha pubescens* on unmown invasive European blackberry (*Rubus anglocandicans*), at three field sites (Belair, Horsnell and Blackwood) in South Australia to measure the impact of infection host performance. Concurrently, we established the parasite on hosts that were mown at two of these sites (Horsnell and Blackwood), to determine the impact of mowing, a commonly used control method, in conjunction with infection by *C. pubescens*.
3. Fruit production, midday quantum yield and electron transport rates of infected *R. anglocandidans* were significantly lower than uninfected plants at only one site, Blackwood. Predawn quantum yield, and foliar nitrogen and phosphorus concentrations of infected plants were significantly lower than uninfected ones across all three sites. Stomatal conductance was negatively affected by infection at one site (Belair). Mowing enhanced parasite impact on host nitrogen concentration at one site (Horsnell), and infection negatively affected host stomatal conductance at the same site, irrespective of whether plants were mown or not.
4. We have demonstrated that this native biocontrol can be artificially established on invasive European blackberry in the field, with negative consequences for its performance. Our results demonstrate the feasibility of implementing native parasitic plants as weed biocontrols to protect biodiversity, and are aligned with the Biotic Resistance hypothesis that invasive species are susceptible and sensitive to enemies native to their newly invaded habitat.

## 1 INTRODUCTION

Invasive plants impact ecosystem quality and displace native flora and fauna, decreasing biodiversity. They also affect water reserves and food production, and promote further invasions (Vilà et al., 2019). Once established, measures of control can be costly, difficult to apply and vary in efficacy (Culliney, 2005; Diagne et al., 2021). Biocontrol is generally the most cost-effective and environmentally sound control method, and is particularly useful for large infestations and sensitive or difficult to access areas (Culliney, 2005). Biocontrols include natural enemies introduced from the invasive species’ native range, or new enemies native to the invaded habitat. Within the framework of invasion theory, two hypotheses describe these different biocontrol mechanisms. The enemy release hypothesis postulates that invasive species are successful because they are released from their native enemies (i.e. classical biocontrol) (Parker et al., 2006). The biotic resistance hypothesis states that invasive species are stifled by encounters with new enemies (i.e. native biocontrol) native to their newly invaded range (Parker et al., 2006). Classical and native biocontrol can both be effective weed management tools (Parker et al., 2006; Clewley et al., 2012), however, native biocontrol is preferred because it usually involves less risk (Verhoeven et al., 2009).

Evidence suggests that native parasitic plants have potential as biocontrol for some of the world’s worst weeds (Těšitel et al., 2020). For example, in Australia, the native hemiparasitic vine *Cassytha pubescens* generally has a strong negative impact on *Ulex europaeus*, one of the world’s 100 worst invasive species (Lowe et al., 2000), and *Cytisus scoparius*, than on native hosts studied (Prider et al., 2009; Cirocco et al. 2016a, 2017; but see Cirocco et al. 2021a). To our knowledge, no studies have explored the deliberate application of native parasites in controlling invasive weeds.

Here we report results from field trials investigating the impact of *C. pubescens* on *Rubus anglocandicans* (the most prevalent species in the *R. fruticosus* agg. in Australia; Evans & Weber, 2003). First, we investigated the impact of *C. pubescens* on the performance of *R. anglocandicans* across three sites. We predicted that *C. pubescens* would negatively affect this major invasive weed as has been reported for other invasive hosts (Prider et al., 2009; Shen et al., 2010; Prider et al., 2011; Cirocco et al., 2016a, 2016b, 2017, 2018, 2020, 2021b). Secondly, we also investigated the impact of mowing, on the *C. pubescens-R*.

*anglocandicans* association. Mowing is a frequently used control for *R. fruticosus* agg., also promoting new shoot growth that is prone to rust infection (Amor et al., 1998), and thus, may make this invasive host more sensitive to *C. pubescens*. To quantify parasite performance and impact on the invasive host, we measured a number of plant traits shown to be affected by *C. pubescens* in other studies, including: light-use efficiency (predawn and midday quantum yield) and electron transport rates (proxy for photosynthesis), which may decline in the host as a result of infection; stomatal conductance, a key indicator of host water stress; stable carbon isotope composition, as long-term indicator of water use efficiency, and nutrient-status which can be adversely affected by parasite removal of resources.

## 2 MATERIALS AND METHODS

### 2.1 Study species

#### Rubus anglocandicans

A. Newton is a perennial shrub (2–3 m in height) with biennial canes armed with prickles (Amor et al., 1998). It is a major invasive weed in many parts of the world and one of the worst weeds in Australia (Parsons & Cuthbertson, 2001). It is difficult to control with conventional methods and has few biocontrol options available (Amor et al., 1998).

#### Cassytha pubescens

R. Br. (Lauraceae) is an Australian native perennial, hemiparasitic vine (Kokubugata et al., 2012). Its stems (0.5–1.5 mm in diameter) coil around and attach to host stems with multiple haustoria (McLuckie, 1924). Being a vine with indeterminate growth, it can infect multiple individuals at any one time and is a generalist parasite commonly infecting perennial shrubby hosts (McLuckie, 1924). *C. pubescens* is known to infect *R. anglocandicans* in the Mt Lofty Ranges of South Australia (pers. obs.).

### 2.2 Study sites and design

Two separate field experiments were conducted in the Mt Lofty Ranges, South Australia. At all field sites, the major invasive host, *R. anglocandicans* was already naturally occurring in dense stands, approximately 1 –2 m tall. First, we established *C. pubescens* on *R. anglocandicans* at three sites. Belair National Park (35°01’97”S, 138°66’61”E), and Horsnell Gully Conservation Park (34°93’29”S, 138°70’26”E), are located within Eucalypt dominated woodland with sclerophyllous understorey. The third site, Blackwood Forest Recreation Park (35°02’88”S, 138°63’15”E), is situated in a *Pinus radiata* plantation. We quantified environmental conditions (light, air temperature and relative humidity) for these sites when physiological measurements were made (Supporting Information Figs S1-S4). Secondly, we also established *C. pubescens* on mown *R. anglocandicans* at Horsnell and Blackwood, but in separate locations from the first experiment. At each site, two 3m × 3m plots were mown (to around 50 cm in height), one plot was left as a control (i.e. uninfected) and in the second we introduced the parasite. This experiment compared uninfected and infected canes in unmown and mown areas at both sites.

The parasite was introduced using the ‘donor plant’ technique (Shen et al., 2010). Briefly, pots containing infected hosts (‘donor plants’) were placed adjacent to *R. anglocandicans* and, over time, attached to host canes and leaf petioles. This was a challenging process because we had to identify sites that were accessible, obtain permission to run the experiments on these sites, have sufficient donor plants, and deploy them effectively amongst dense patches of extremely prickly host plants. We also needed to visit sites at least twice a week to maintain and water donor plants (and to also remove their flowers) to keep them alive long enough for the parasite to establish on the target hosts. The infection process was initiated late June-early July 2018, and the parasite was established on *R. anglocandicans* by Dec 2018 (i.e. treatment imposed), thus, ‘donor plants’ had to be maintained and watered for at least five months. We considered a single cane as a replicate, as the impact of *C. pubescens* is localised to infected *R. anglocandicans* canes (McDowell, 2002). Measurements were made on host canes either without (uninfected cane) or with the parasite (infected cane), in January–February 2019 (data not shown), and in March– April 2019. Replicate number is shown in figure captions.

### 2.3 Fruit and prickle production

Fruit (per cane) and prickles (from cane tip to 30cm below) were counted on uninfected and infected canes of *R. anglocandicans* 63 days after treatment (DAT; i.e. parasite establishment).

### 2.4 Chlorophyll fluorescence, stomatal conductance, δ^13^C and foliar nutrients

Host and parasite predawn (*F*_v_/*F*_m_) and midday quantum yields (Φ_PSII_) and electron transport rates (ETR) were measured 117–124 DAT, with a portable, chlorophyll fluorometer (MINI-PAM and 2030–B leaf clip, Walz, Effeltrich, Germany). Φ_PSII_ and midday ETR measurements were made on sunny or mostly sunny days (13:00-15:45), under natural light, or if light was low, using the internal light from the MINI-PAM (Supporting Information Figs S1–S2). ΦP_SII_ and ETR measurements are sensitive to light, so we ensured light levels were similar for all measurements: mean PPFD (μmol m^-2^ s^-1^) was 1554 ± 8 (*n* = 134).

Host stomatal conductance (g_s_) was measured 125–132 DAT, with a porometer (SC-1, Decagon Devices, Inc. Washington). Despite clear days, plants at the sites were exposed to sunlight at different times of the day due to inherent site differences in elevation, aspect and canopy cover in combination with the lower sun angle at this time of year (Autumn, southern hemisphere). Nevertheless, plants were only measured after they had been exposed to full sunlight for at least 30 min and to account for these differences in timing of measurements, g_s_ was compared within site (Supporting Information Figs S3–S4).

A single leaf from uninfected and infected canes was collected 125–132 DAT, oven-dried at 60°C for 7 days, then finely ground for the following analyses. Foliar carbon isotope composition (δ^13^C) and nitrogen concentration [N] of *R. anglocandicans* were quantified by mass spectrometer (IsoPrime, GV Instruments, Manchester, UK) and elemental analyser (Elementar Isotope CUBE, Elementar Analysensysteme, Hanau, Germany). Foliar concentrations of phosphorus [P], sodium [Na] and iron [Fe] were determined using inductively coupled plasma spectroscopy (Cuming Smith British Petroleum Soil and Plant Laboratory, Western Australia).

### 2.5 Statistical analyses

For the first experiment we examined the interactive effects of infection and site on performance of unmown *R. anglocandicans* using two-way ANOVA and the main effect of site on performance of *C. pubescens* with one-way ANOVA. For the second experiment we examined the interactive effects of infection, site and mowing on performance of *R. anglocandicans* using three-way ANOVA and site and mowing effects on parasite performance of *C. pubescens* with two-way ANOVA. In all analyses, we have included sites as a fixed (not random) factor, because as pointed out earlier, we were limited in the number of donor plants/sites available. Thus, we emphasise that the results pertain to these sites and inferences beyond our experimental conditions should be made with caution. Nevertheless, sites/mown plots were replicated, hence avoiding pseudoreplication and ensuring robust results. Significant interactions were subjected to Tukey HSD. When interactions were not detected, main effects of infection, site, or mowing were considered valid. As mentioned, to account for light differences among sites when conducting host g_s_ measurements, we, examined the effect of infection, and infection and mowing within sites for the first and second experiments, respectively. Model assumptions were met, in some cases after transformation where stated, data were analysed using R (R Development Core Team, 2016) and *α* = 0.05 (Type I error rate).

## 3 RESULTS

### 3.1 Fruit and prickles of *R. anglocandicans*

There was a significant infection × site interaction for fruit production of *R. anglocandicans* (Table 1). At Blackwood, infected canes had 55% fewer fruit than uninfected ones, while infection did not significantly affect fruit production at the other two sites (Figure 1a). There were main effects of infection and site for number of prickles on *R. anglocandicans* (Table 1; no interaction: Supporting Information Figure S5a). There were 13% fewer prickles on infected canes relative to uninfected ones (Figure S5b) and prickle number was significantly higher at Blackwood relative to the other two sites (Figure S5c).

**TABLE 1.**
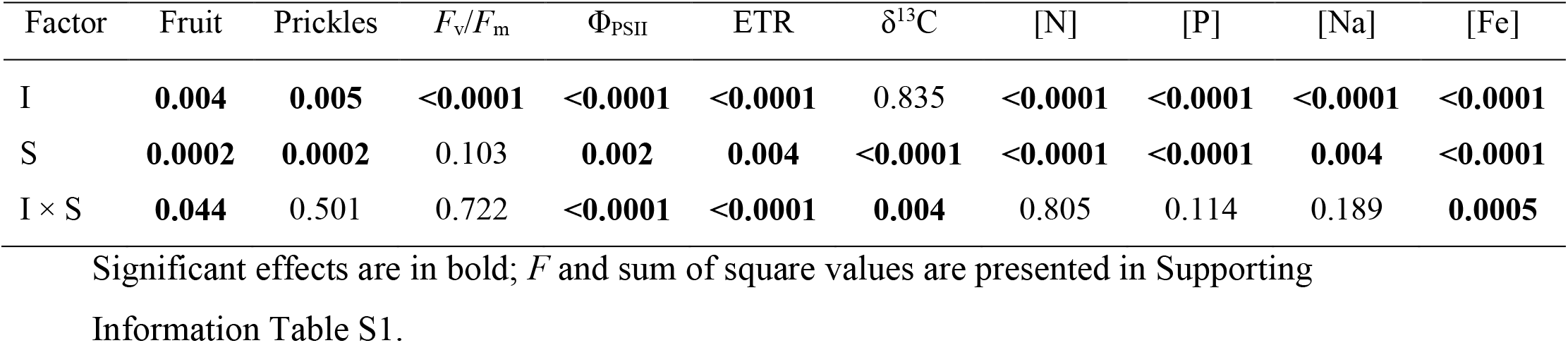
Two-way ANOVA results (*p*-values) for the effects of infection with *Cassytha pubescens* (I) and site (S) on number of fruits, number of prickles, predawn and midday quantum yield (*F*_v_/*F*_m_ and Φ_PSII_), midday electron transport rates (ETR), foliar carbon isotope composition (δ^13^C), nitrogen [N], phosphorus [P], sodium [Na] and iron concentration [Fe] of *Rubus anglocandicans*

| Factor | Fruit | Prickles | $F_v/F_m$ | $\Phi_{PSII}$ | ETR | $\delta^{13}C$ | [N] | [P] | [Na] | [Fe] |
| --- | --- | --- | --- | --- | --- | --- | --- | --- | --- | --- |
| I | **0.004** | **0.005** | **<0.0001** | **<0.0001** | **<0.0001** | 0.835 | **<0.0001** | **<0.0001** | **<0.0001** | **<0.0001** |
| S | **0.0002** | **0.0002** | 0.103 | **0.002** | **0.004** | **<0.0001** | **<0.0001** | **<0.0001** | **0.004** | **<0.0001** |
| I × S | **0.044** | 0.501 | 0.722 | **<0.0001** | **<0.0001** | **0.004** | 0.805 | 0.114 | 0.189 | **0.0005** |

Significant effects are in bold; *F* and sum of square values are presented in Supporting Information Table S1.

**FIGURE 1.**
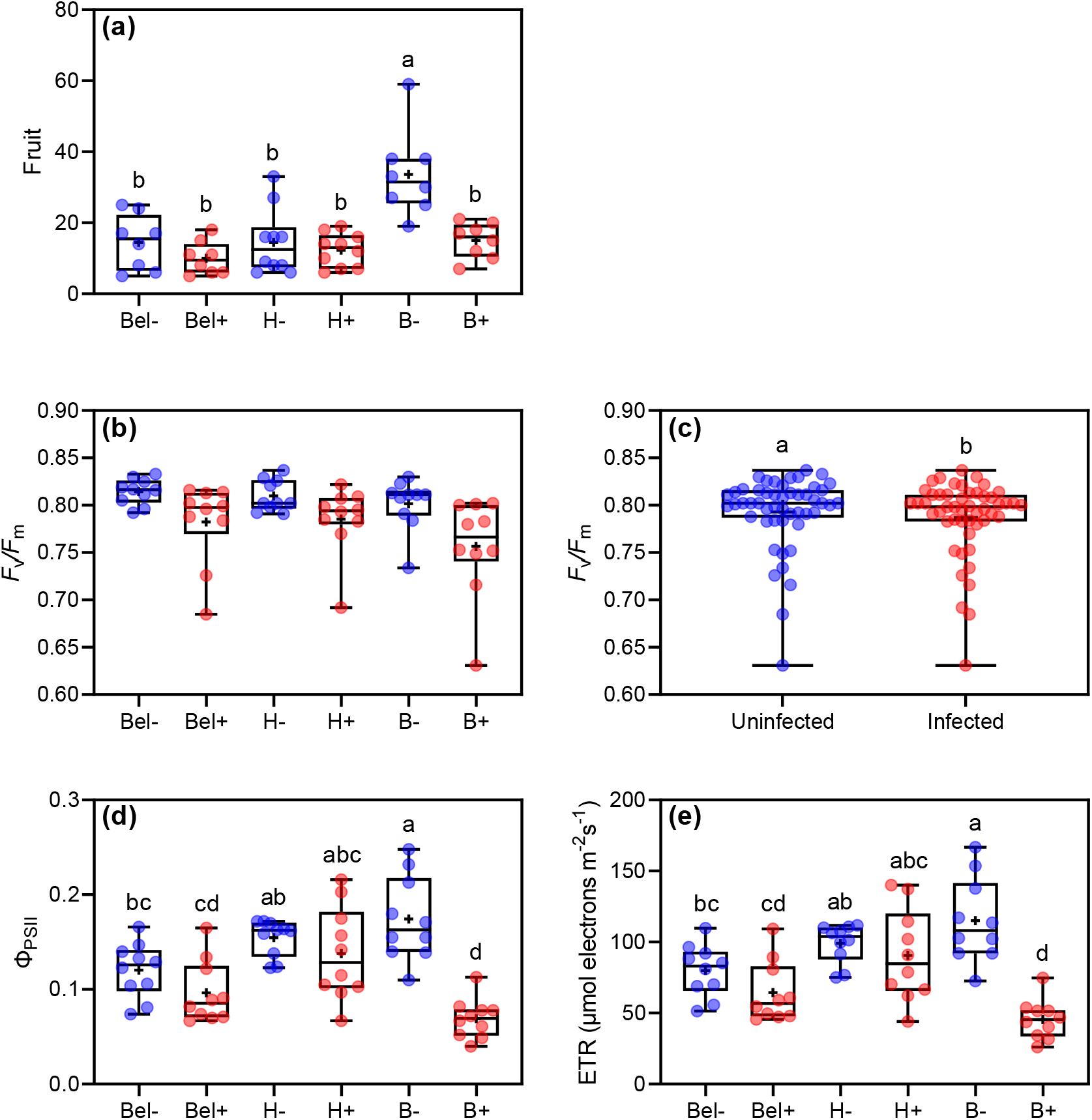
(a) Number of fruit (per cane), (b and d), predawn (*F*_v_/*F*_m_) and midday (Φ_PSII_) quantum yield, and (e) midday electron transport rates (ETR) of *Rubus anglocandicans*, when uninfected (–) or infected (+) with *Cassytha pubescens* at Belair (Bel–, Bel+), Horsnell (H–, H+) and Blackwood (B–, B+), respectively. (c) Main effect of infection on host *F*_v_/*F*_m_. All data points, median, percentile lines and mean (+ within box) are displayed, different letters indicate significant differences: (a) *n* = 8–10, (b, d, e) *n* = 10 and (c) *n* = 20

For the mowing experiment, as virtually no fruit was produced in the mown area at Blackwood, we only report on fruit and prickle production at Horsnell. No significant infection × mowing effect (*F*_1,24_ = 0.444, *p* = 0.512) or main effect of infection (*F*_1,24_ = 2.06, *p* = 0.164) or mowing (*F*_1,24_ = 0.381, *p* = 0.543) were found for host fruit production at Horsnell (Figure 2a). There was a significant infection × mowing effect for prickle number (*F*_1,24_ = 4.68, *p* = 0.041). Number of prickles on infected plants was 20% lower than that of uninfected plants in the unmown area while infection had no effect on this variable in the mown plots (Supporting Information Figure S6a).

**FIGURE 2.**
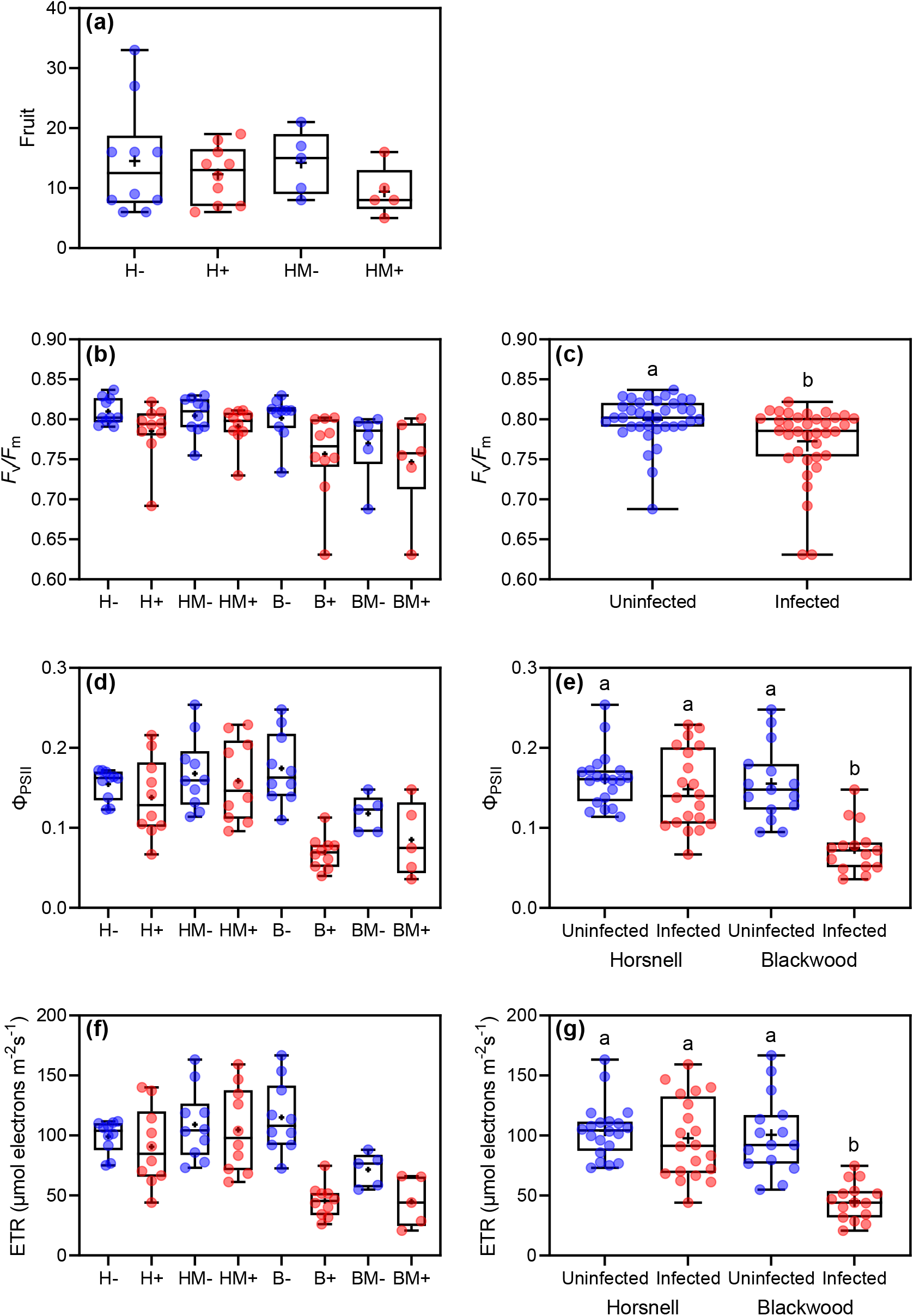
(a) Number of fruit (per cane), (b and d), predawn (*F*_v_/*F*_m_) and midday (Φ_PSII_) quantum yield, and (f) midday electron transport rates (ETR) of *Rubus anglocandicans*, when unmown or mown (m) and uninfected (-) or infected (+) with *Cassytha pubescens* at Horsnell (unmown: H-, H+, mown: HM-, HM+) and Blackwood (unmown: B-, B+, mown: BM-, BM+), respectively. (c) Main effect of infection on host *F*_v_/*F*_m_. Infection × site interaction on host (e) Φ_PSII_ and (g) ETR. All data points, median, percentile lines and mean (+ within box) are displayed, different letters indicate significant differences: (a, d, f) *n* = 5-10, (b) *n* = 6-10, (c) *n* = 36, (e) *n* = 20 and (g) *n* = 15-20 (Blackwood and Horsnell, respectively)

### 3.2 Host and parasite photosynthetic performance

There was a main effect of infection on *F*_v_/*F*_m_ of *R. anglocandicans* (Table 1; no interaction: Figure 1b). *F*_v_/*F*_m_ of infected plants was 4% lower than that of uninfected plants (Figure 1c). A significant infection × site interaction was found for Φ_PSII_ and ETR (Table 1). At Blackwood, Φ_PSII_ and ETR of infected plants were 60% and 55% lower than uninfected plants, respectively, while infection did not affect these variables at the other two sites (Figure 1d,e).

*F*_v_/*F*_m_ of *C. pubescens* was significantly lower at Horsnell relative to Belair and intermediate at Blackwood (*p* = 0.020; Figure 3a; Table S3). Parasite Φ_PSII_ (*p* = 0.003) and ETR (*p* = 0.0008) were significantly higher at Horsnell than at the other two sites (Figure 3b,c; Table S3).

**FIGURE 3.**
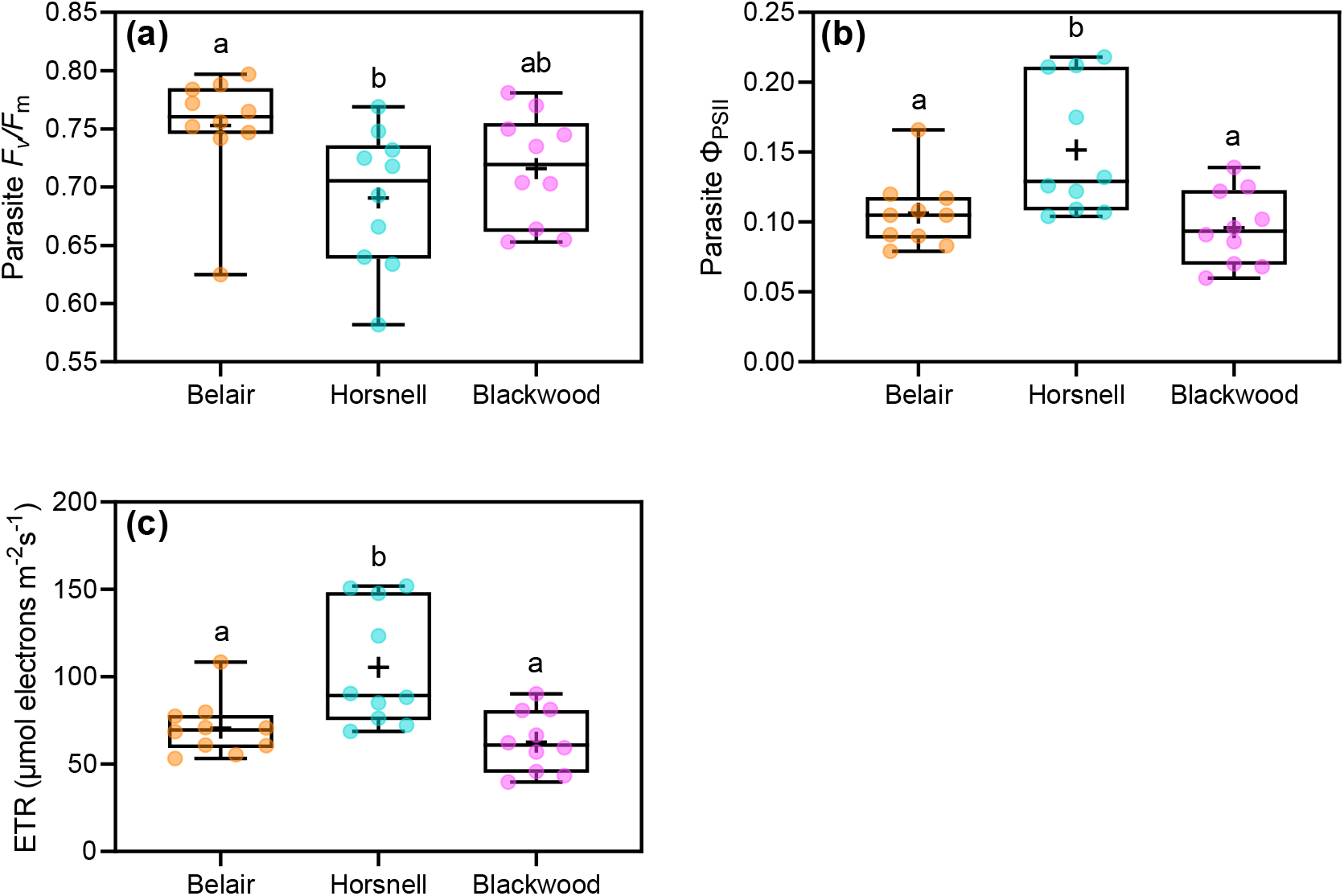
(a) Predawn (*F*_v_/*F*_m_) and (b) midday quantum yield (Φ_PSII_), and (c) midday electron transport rates (ETR) of *Cassytha pubescens* (infecting *R. anglocandicans*) at Belair, Horsnell, and Blackwood. All data points, median, percentile lines and mean (+ within box) are displayed, different letters indicate significant differences: (a, b, c) *n* = 10

For the mowing experiment, there was a main effect of infection on *F*_v_/*F*_m_ of *R. anglocandicans* (Table 2; no three-way interaction: Figure 2b). *F*_v_/*F*_m_ of infected plants was 3% lower than that of uninfected ones (Figure 2c). There was also a main effect of site on this variable; *F*_v_/*F*_m_ of *R. anglocandicans* was 3% higher at Horsnell relative to that at Blackwood (Supporting Information Figure S6b). An infection × site interaction was found for Φ_PSII_ and ETR of *R. anglocandicans* (Table 2; no three-way interaction: Figure 2d,f,). At Blackwood, Φ_PSII_ and ETR of infected plants were, respectively, 52% and 55% lower than uninfected plants, whereas infection had no effect on this variable at Horsnell (Figure 2e,g). A site × mowing interaction was found for host ETR, which was significantly lower for mown plants at Blackwood but not at Horsnell (Table 2; Supporting Information Figure S6c).

**TABLE 2.** Three-way ANOVA results (*p*-values) for the effects of infection with *Cassytha pubescens* (I), site (S) and mowing (M) on predawn and midday quantum yield (*F*_v_/*F*_m_ and Φ_PSII_), midday electron transport rates (ETR), foliar carbon isotope composition (δ^13^C), nitrogen [N], phosphorus [P], sodium [Na] and iron concentration [Fe] of *Rubus anglocandicans*

| | $F_v/F_m$ | $\Phi_{PSII}$ | ETR | $\delta^{13}C$ | [N] | [P] | [Na] | [Fe] |
| --- | --- | --- | --- | --- | --- | --- | --- | --- |
| I | **0.0005** | **<0.0001** | **<0.0001** | **0.050** | **<0.0001** | **0.003** | **<0.0001** | **<0.0001** |
| S | **0.001** | **<0.0001** | **<0.0001** | **<0.0001** | **<0.0001** | **<0.0001** | **<0.0001** | **<0.0001** |
| I × S | 0.551 | **0.0005** | **0.0001** | **0.025** | 0.242 | 0.168 | **0.045** | **0.0004** |
| M | 0.227 | 0.844 | 0.710 | **<0.0001** | **<0.0001** | **0.0005** | **0.002** | **0.0004** |
| I × M | 0.262 | 0.080 | 0.131 | 0.962 | 0.362 | 0.791 | 0.535 | 0.182 |
| S × M | 0.205 | 0.089 | **0.012** | 0.124 | 0.262 | 0.165 | **0.0008** | 0.372 |
| I × S × M | 0.600 | 0.133 | 0.186 | 0.236 | **0.045** | 0.690 | 0.594 | 0.939 |

Significant effects are in bold; *F* and sum of square values are presented in Supporting Information Table S2.

No significant site × mowing interaction (*p* = 0.207) or main effects of site (*p* = 0.675) or mowing (*p* = 0.118) were detected for *F*_v_/*F*_m_ of *C. pubescens* (Figure 4a; Table S3). However, there was a site × mowing interaction, detected for parasite Φ_PSII_ (*p* = 0.002) and ETR (*p* = 0.0006), which decreased by 40% and 49%, respectively, in response to mowing at Blackwood but were unaffected by mowing at Horsnell (Figure 4b,c; Table S3).

**FIGURE 4.**
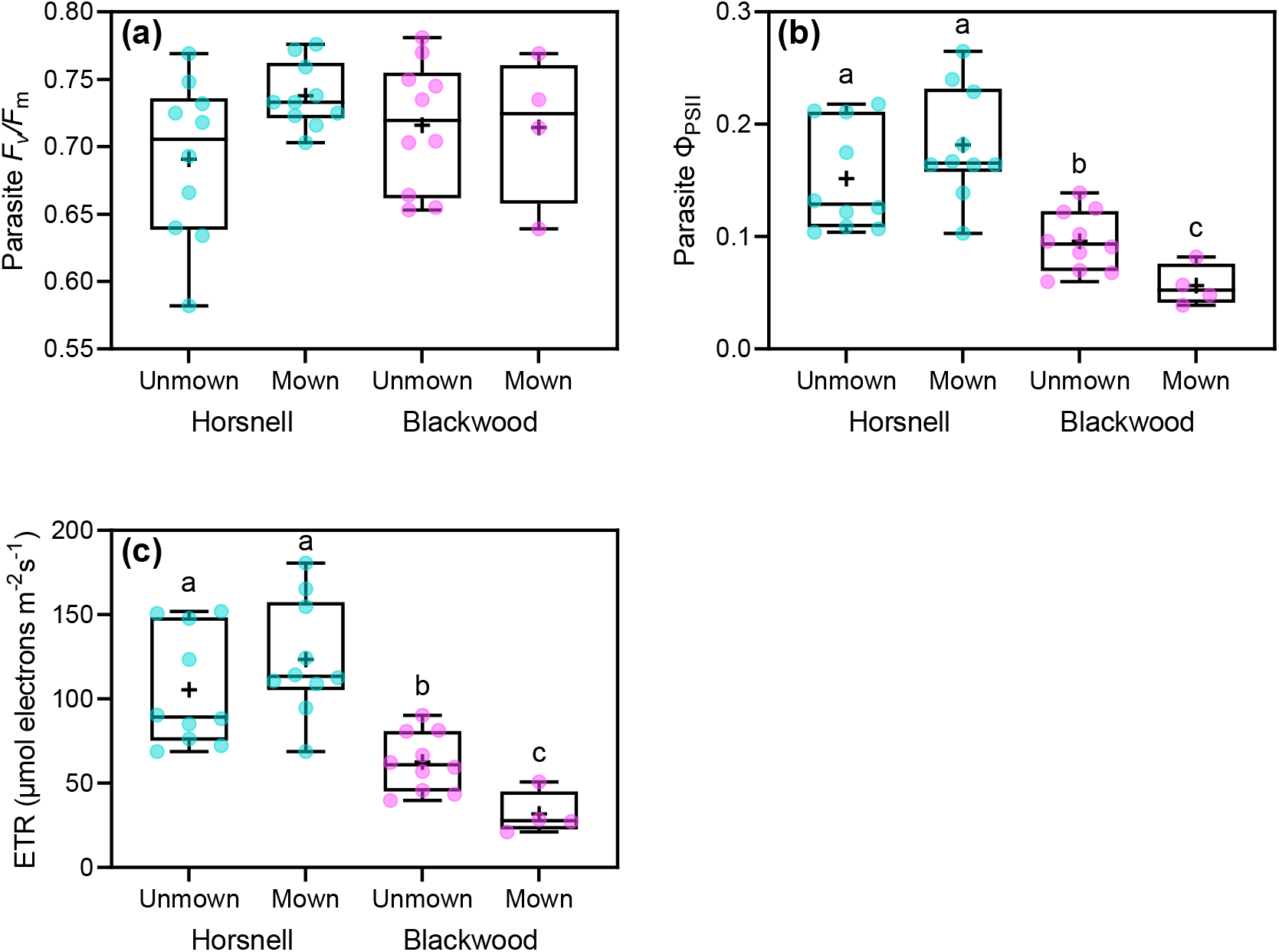
(a) Predawn (*F*_v_/*F*_m_) and (b) midday quantum yield (Φ_PSII_), and (c) midday electron transport rates (ETR) of *Cassytha pubescens* (infecting unmown or mown *R. anglocandicans*) at Horsnell, and Blackwood. All data points, median, percentile lines and mean (+ within box) are displayed, different letters indicate significant differences: (a, b, c) *n* = 10 (except *n* = 4 for Mown plants at Blackwood)

### 3.3 *R. anglocandicans* g_s_, δ^13^C and nutrients

There was a significant negative effect of infection on g_s_ of *R. anglocandicans* at Belair (*p* = 0.0009), but not at Horsnell (*p* = 0.173), or Blackwood (*p* = 0.288) (Figure 5a,b,c; Table S4). At Belair, infected plants had 46% lower g_s_ than uninfected ones (Figure 5a).

**FIGURE 5.**
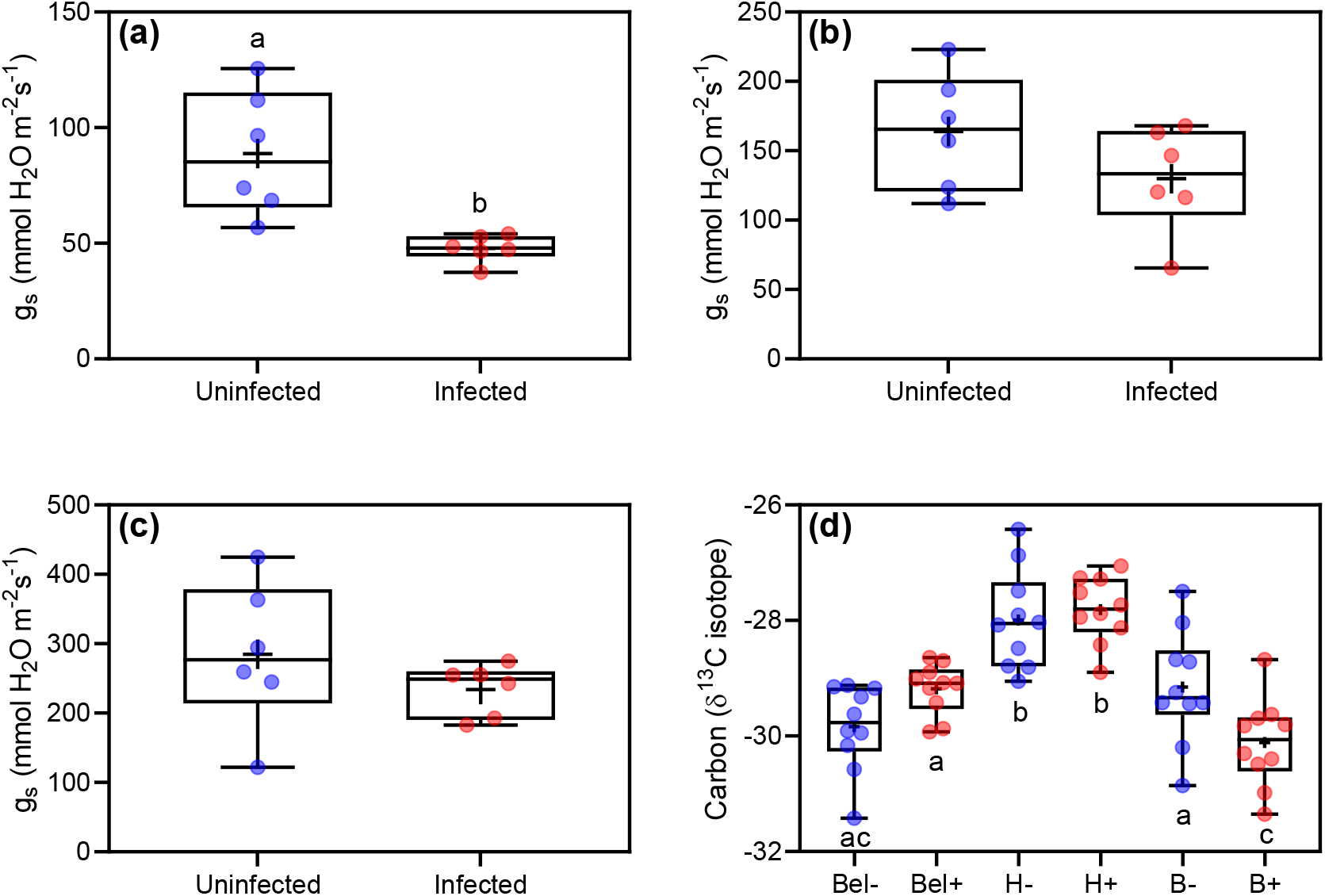
Stomatal conductance (g_s_) of *Rubus anglocandicans* either uninfected (–) or infected (+) with *Cassytha pubescens* at (a) Belair, (b) Horsnell and (c) Blackwood. (d) Carbon isotope composition of *Rubus anglocandicans*, when uninfected (–) or infected (+) with *Cassytha pubescens* at Belair (Bel–, Bel+), Horsnell (H–, H+) and Blackwood (B–, B+), respectively. All data points, median, percentile lines and mean (+ within box) are displayed, different letters indicate significant differences and (a, b, c) *n* = 6 and (d) *n* = 10

For the mowing experiment, there was no infection × mowing interaction on g_s_ of *R. anglocandicans* at either Horsnell (*p* = 0.174) or Blackwood (*p* = 0.680) (Figure 6a,c; Table S4). A main effect of infection was found at Horsnell (*p* = 0.004) but not at Blackwood (*p* = 0.233) (Table S4). Infected plants at Horsnell had 29% lower g_s_ relative to uninfected plants (Figure 6b). A main effect of mowing was also found for host g_s_ at Horsnell (*p* = 0.006) and Blackwood (*p* = 0.002) (Table S4). At Horsnell, g_s_ of unmown plants was 28% lower than that of mown plants (Supporting Information Figure S7a). At Blackwood, g_s_ of unmown plants was 47% higher than mown plants (Supporting Information Figure S7b).

**FIGURE 6.**
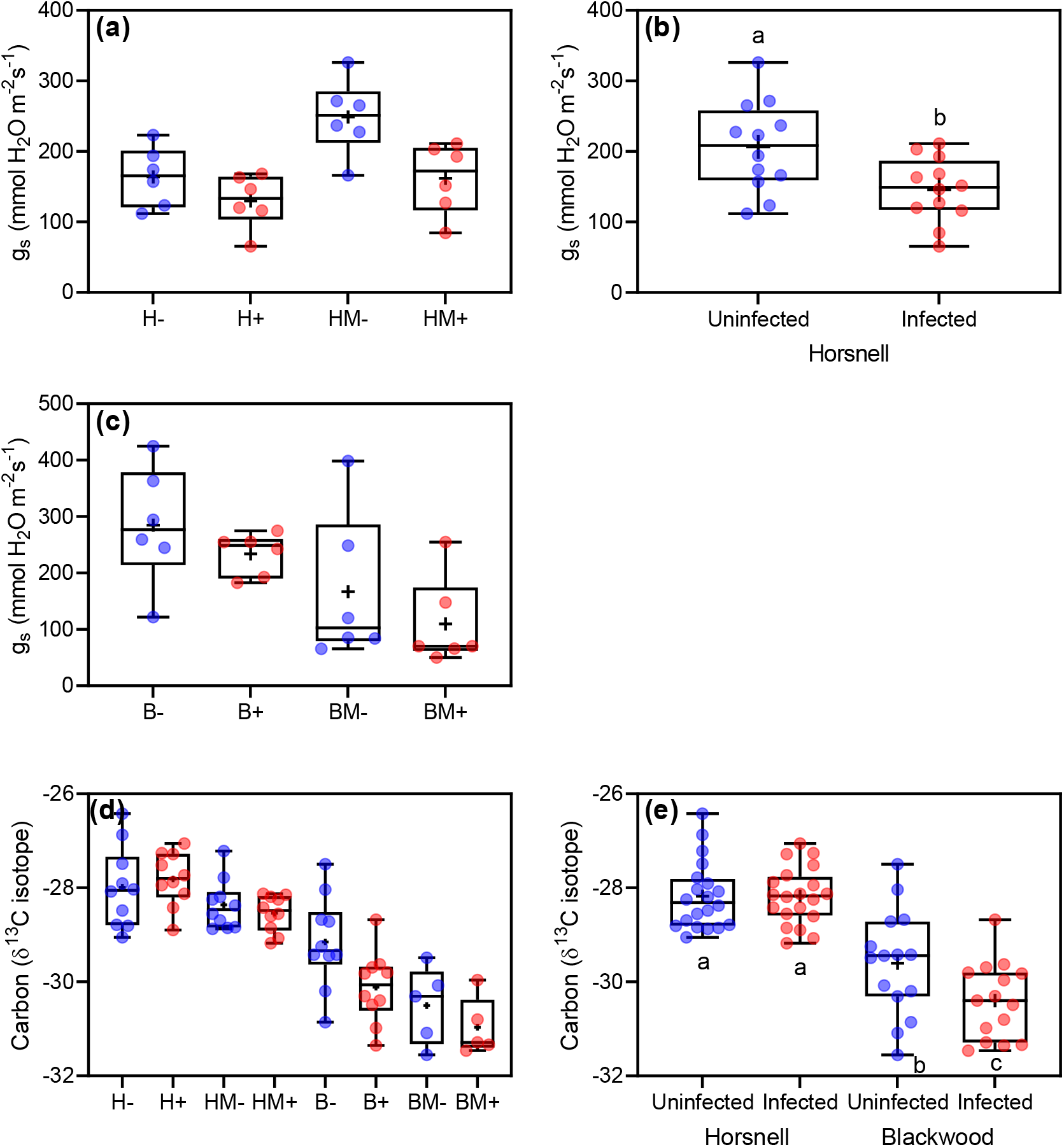
Stomatal conductance (g_s_) of *Rubus anglocandicans*, when unmown or mown (m) and uninfected (-) or infected (+) with *Cassytha pubescens* at (a) Horsnell (unmown: H-, H+, mown: HM-, HM+) and (c) Blackwood (unmown: B-, B+, mown: BM-, BM+), respectively. (b) Main effect of infection on host g_s_ at Horsnell. (d) Carbon isotope composition (δ^13^C) of *R. anglocandicans* and (e) Infection × site effect on host δ^13^C. All data points, median, percentile lines and mean (+ within box) are displayed, different letters indicate significant differences: (a, c), *n* = 6, (b) *n* = 12, (d) *n* = 10 (except *n* = 4 for all Mown plants at Blackwood) and (e) *n* = 15-20 (Blackwood and Horsnell, respectively)

There was an infection × site interaction for δ^13^C *of R. anglocandicans* (Table 1). δ^13^C of infected plants was significantly lower than that of uninfected plants at Blackwood while no significant effect was detected at Horsnell or Belair (Figure 5d).

For the mowing experiment, there was an infection × site interaction for leaf δ^13^C of *R. anglocandicans* (Table 2; no three-way interaction: Figure 6d). δ^13^C of infected plants was significantly lower than that of uninfected plants at Blackwood while no significant effect was detected at Horsnell (Figure 6e). A main effect of mowing was also found for host δ^13^C (Table 2). δ^13^C of mown plants was significantly lower relative to that of unmown ones (Supporting Information Figure S7c).

There was a main effect of infection on foliar [N], [P] and [Na] of *R. anglocandicans* (Tables 1 and 3; no three-way interactions: Figure 7a,c,e). Nitrogen and [P] were 15% and 14% lower, for infected than uninfected plants, respectively, while infection increased host [Na] by 47% (Figure 7b,d,f). There was a main effect of site on [N] and [Na] of *R. anglocandicans* (Table 2). At Horsnell, [N] and [Na] were 12.5% and 23% lower, respectively, than those at the other two sites (Supporting Information Figure S8a,c). Host [P] at Horsnell and Belair was 43% and 25% lower, respectively, than that at Blackwood (Supporting Information Figure S8b). An infection × site interaction was detected for [Fe] (Table 1). Foliar [Fe] of infected plants was *c*. 40% higher than that of uninfected plants at Belair and Blackwood, whereas infection had no effect on this variable at Horsnell (Figure 7g).

**FIGURE 7.**
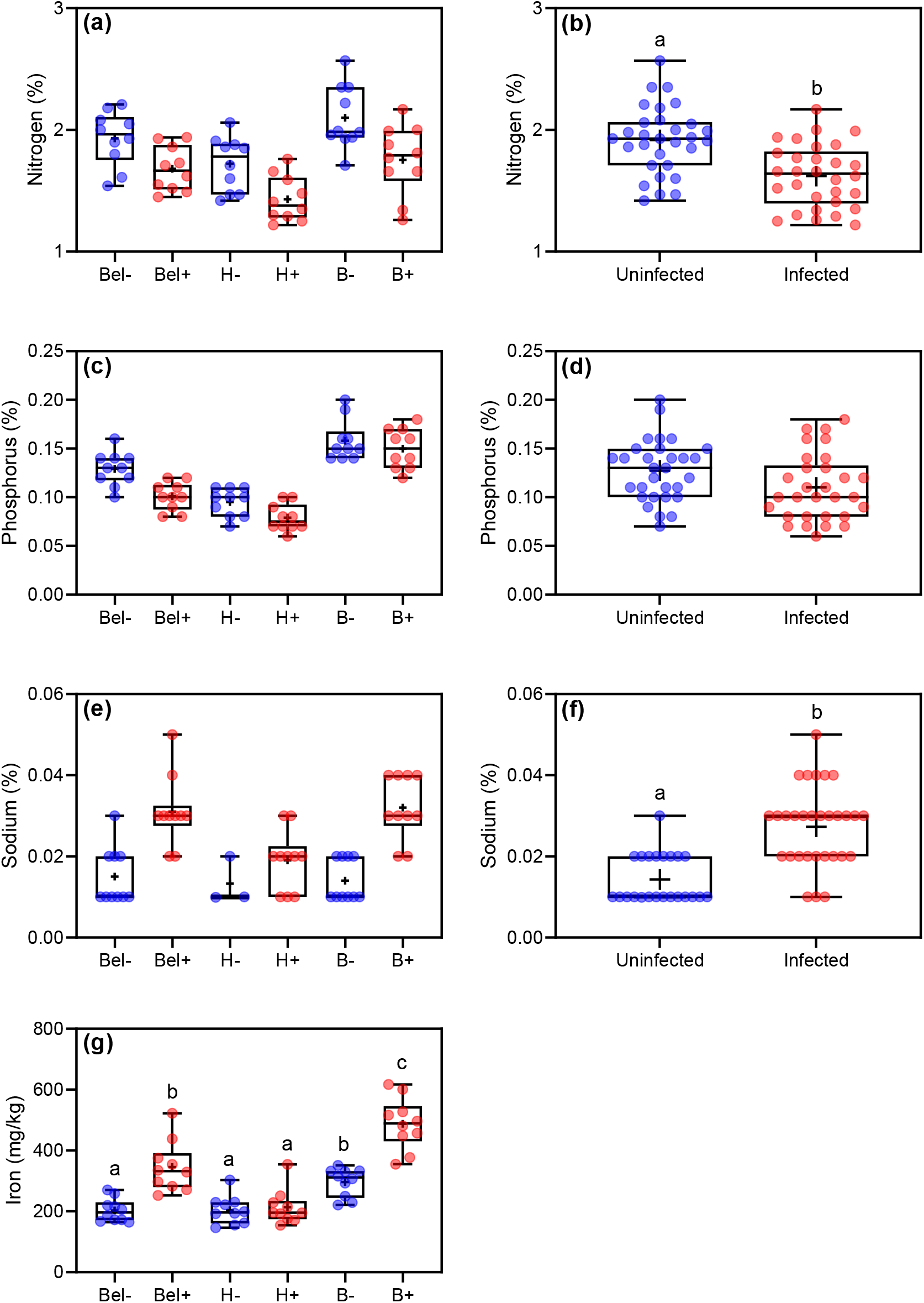
Leaf (a) nitrogen, (c) phosphorus, (e) sodium and (g) iron concentration of *Rubus anglocandicans*, when uninfected (–) or infected (+) with *Cassytha pubescens* at Belair (Bel–, Bel+), Horsnell (H–, H+) and Blackwood (B–, B+), respectively. (c) Main effect of infection on host (b) nitrogen, (d) phosphorus and (f) sodium concentration. All data points, median, percentile lines and mean (+ within box) are displayed, different letters indicate significant differences: (a, c, g) *n* = 10, (b, d) *n* = 30, (e) *n* = 10 (except *n* = 3 for uninfected plants at Horsnell: H-, because sodium levels were too low for the instrument to detect) and (f) *n* = 23-30 (uninfected infected plants, respectively)

For the mowing experiment, an infection × site × mowing interaction was detected for foliar [N] of *R. anglocandicans* (Table 2). Infection negatively affected host [N] when plants were mown at Horsnell (by 25%) and when plants were unmown at Blackwood (by 16%) (Figure 8a). There was a main effect of infection on host [P] (Table 2; no three-way interaction: Figure 8b). Foliar [P] of infected plants was 9% lower than that of uninfected plants (Figure 8c). There were also main effects of site and mowing on host [P], which was 47% lower at Horsnell relative to Blackwood, and 20% lower in mown plots compared with unmown plots (Supporting Information Figure S9a,b). There was an infection × site interaction found for host [Na] and [Fe] (Table 2; no three-way interaction: Figure 8d,f). At Blackwood [Na] and [Fe] of infected plants were 52% and 44%, higher than uninfected plants, respectively, but infection induced non-significant increases in host [Na] and [Fe] at Horsnell (Figure 8e,g). There was a site × mowing effect detected for host [Na] (Table 2). Mowing resulted in host [Na] being significantly higher at Blackwood, but not at Horsnell (Supporting Information Figure S9c). There was a main effect of mowing on host [Fe] (Table 2). Host [Fe] significantly increased when plants were mown (Supporting Information Figure S9d).

**FIGURE 8.**
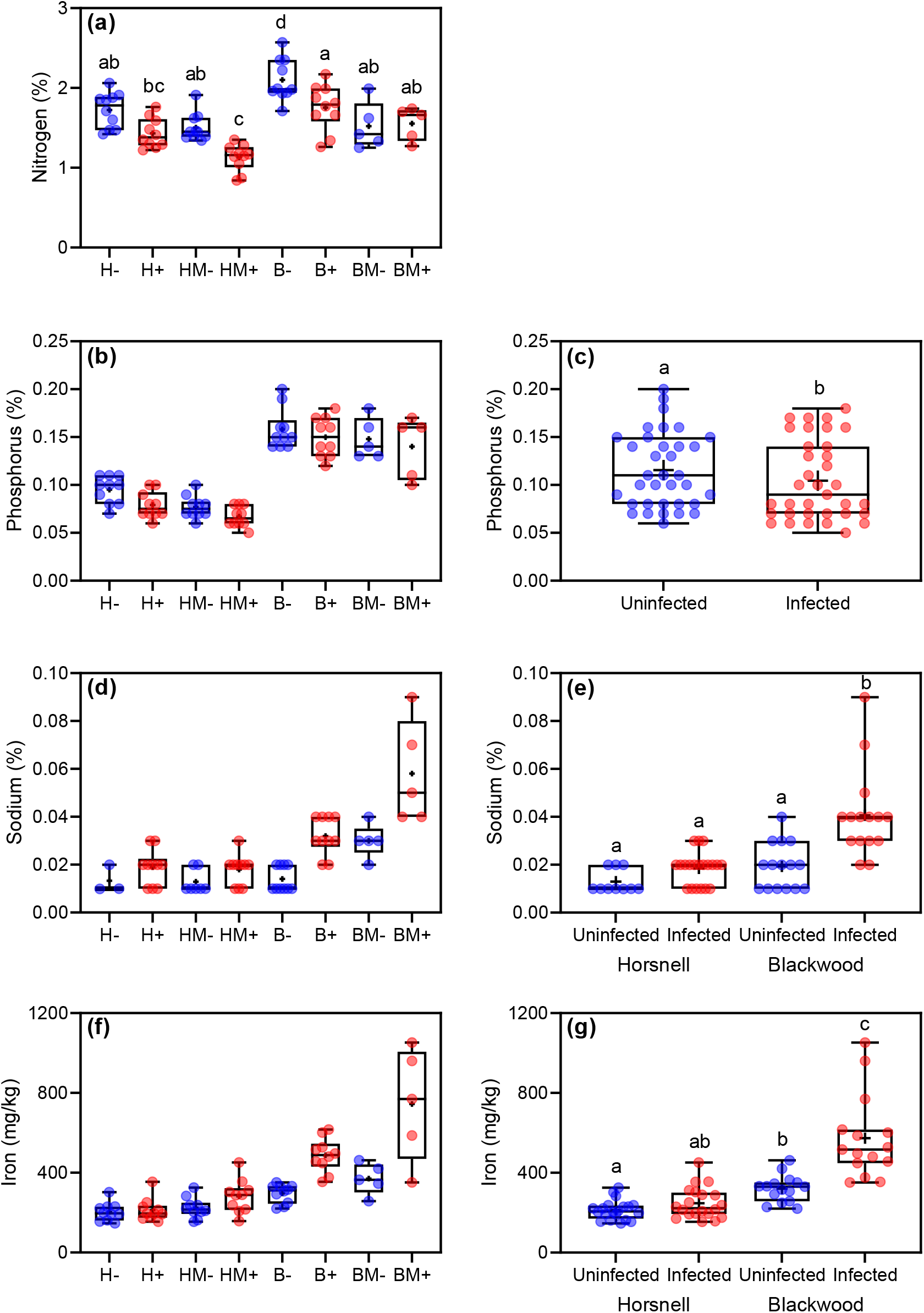
Leaf (a) nitrogen, (b) phosphorus, (d) sodium and (f) iron concentration of *Rubus anglocandicans*, when unmown or mown (m) and uninfected (-) or infected (+) with *Cassytha pubescens* at (a) Horsnell (unmown: H-, H+, mown: HM-, HM+) and (c) Blackwood (unmown: B-, B+, mown: BM-, BM+), respectively. (c) Main effect of infection on host phosphorus. Infection × site interaction on host (e) sodium and (g) iron. All data points, median, percentile lines and mean (+ within box) are displayed, different letters indicate significant differences: (a, b, f) *n* = 10 (except *n* = 5 for BM- and BM+), (c) *n* = 35, (d) *n* = 3-10, (e) *n* = 10-20 and (g) *n* = 15-20

## 4 DISCUSSION

Infection had a significant negative effect on performance of *R. anglocandicans* only four months after parasite establishment, including on fruit production and *F*_v_/*F*_m_; although the effect varied with site. It should be noted that the parasite also had a significant negative effect on host *F*_v_/*F*_m_ and g_s_ across the three sites, just one month after establishment (data not shown). Significant interactions between mowing and infection were only found at one site, where mowing resulted in lower N in infected plants relative to uninfected plants. This demonstrates that *C. pubescens* negatively impacts invasive *R. anglocandicans*, similar to previous reports for other invasive hosts (Prider et al., 2009; Shen et al., 2010; Cirocco et al., 2016b, 2018, 2020, 2021b), but that mowing does little to enhance the impact of the parasite on this host. In contrast, a study by Těšitel et al. (2017) found that the native root hemiparasite *Rhinanthus alectorolophus* more negatively affected the biomass of the expansive native grass *Calamagrostis epigejos* when mowing intensity was increased (no unmown treatment included).

The negative impact on fruit production we observed is important because of its potential effect on host fitness, and dispersal to other locations. Fruit and seed production of the invasive host *Cytisus scoparius* were also negatively impacted by *C. pubescens* (Prider et al., 2011). Similarly, in China, three invasive hosts produced significantly fewer flower stalks when infected with *Cuscuta* spp. (Yu et al., 2008, 2011). Here, the greater negative impact of infection on fruit production at Blackwood, relative to the other two sites may be due to stronger infection effects on host photosynthesis found for this site (Figure 1d,e).

Importantly, infection negatively impacted *F*_v_/*F*_m_ of *R. anglocandidans*. Similarly, *C. pubescens* had a significant negative effect on *F*_v_/*F*_m_ of the major invasive hosts, *Ulex europaeus* and *Cytisus scoparius* (Shen et al. 2010; Cirocco et al. 2016b, 2020, 2021b; but see Prider et al. 2009). In contrast, *C. pubescens* had no impact on *F*_v_/*F*_m_ of the native host *Leptospermum myrsinoides* (Prider et al. 2009; Cirocco et al. 2015). Cirocco et al. (2021a) found that *C. pubescens* only affected *F*_v_/*F*_m_ of the native host, *Acacia paradoxa*, in low phosphorus conditions. Other studies in China, have also found that *Cuscuta* spp. negatively affect the light-use efficiency of the major invasive host, *Mikania micrantha* (Shen et al. 2007, 2013; Le et al. 2015). Here, the negative effect of infection on host *F*_v_/*F*_m_ indicates that infected plants were experiencing chronic photoinhibition (Demmig-Adams and Adams, 2006), a condition that could result in longer-term impacts on growth and persistence of these invasive hosts. This may have eventuated from infected plants being exposed to excess light for prolonged periods as a result of host photosynthesis declining at a constant or increasing PFD (Demmig-Adams and Adams, 1992). The parasite negatively affected host ETR at all sites, but only significantly so at Blackwood where the lowest *F*_v_/*F*_m_ values were recorded.

In our study, parasite-induced decreases in g_s_, [N] and [P] levels may underpin negative parasite effects on host photosynthetic performance (as indicated by *F*_v_/*F*_m_) (Evans, 1989; Rychter & Rao, 2005). *Cassytha pubescens* and the native *Cuscuta australis* have also been found to negatively affect g_s_ of other invasive hosts (Shen et al., 2010; Le et al., 2015). Perhaps the most studied hemiparasite, *Striga*, also adversely affects host stomatal conductance (Watling & Press, 1997; Taylor & Seel, 1998). Lower g_s_ could also result in higher leaf temperatures, which along with CO_2_ limitation would impact host photosynthesis. Lower [N] and [P] will both affect photosynthesis, although impacts of *C. pubescens* on invasive host [N] are known to vary (e.g. Cirocco et al., 2016b, 2018, 2021b; but also see Cirocco et al., 2016a; 2017, 2020). In contrast, *C. pubescens* has been found to have no impact on [N] of native hosts (Cirocco et al., 2016a, 2017, 2021a). In China, *Cuscuta* spp. were also found to negatively impact [N] and [P] of three invasive species (Yu et al., 2009, 2011). Here, the negative effect of infection on host g_s_, [N] and [P] are likely due to the removal of resources by the parasite.

Interestingly, we also found that infection resulted in significant increases in foliar [Na] of *R. anglocandicans*. Other parasitic plants capable of photosynthesis (hemiparasites) have been found to have no effect (Struthers et al., 1986; Tennakoon & Pate, 1996; Lo Gullo et al., 2012) or significantly decrease host foliar [Na] levels (Mutlu et al., 2016; Al-Rowaily et al., 2020). The fact that infected plants were especially enriched in [Na] at Blackwood (Figure 8e) in tandem with no significant parasite-induced decrease in g_s_ at this site (Figures 5c, 6c), is consistent with infected plants lowering their water potential. This is plausible considering that the significantly lower δ^13^C of infected plants at Blackwood (Figures 5d, 6e) indicates that they were being less conservative in their water-use. More profligate water-use leading to lower water potentials would facilitate water uptake and help offset water loss to the parasite, while also making it more difficult for *C. pubescens* to extract resources (Cirocco et al. 2016b). Infected plants at Blackwood suffered most from infection (i.e. lower nitrogen, Figure 8a, and photosynthesis), and the lowering of water potential by the host may have been triggered by this.

Increased uptake of sodium by infected plants may result in charge imbalance in host root cells causing release of protons, and a more acidified rhizosphere (Haynes, 1990). The latter would lead to increased mobility of Fe in the soil and explain why infected plants at Belair and Blackwood had significantly higher foliar [Fe] than uninfected plants. Cirocco et al. (2018) also found that infection with *C. pubescens* resulted in significant enrichment of foliar [Fe] of *U europaeus* across three field sites (Cirocco et al., 2018). Cirocco et al. (2020) found that *C. pubescens* induced significant increases in [Fe] of small but not large *U. europaeus*. On the other hand, the mistletoe *Viscum album* significantly decreased foliar [Fe] of Scots pine (Mutlu et al., 2016), while five different mistletoes (Tennakoon & Pate, 1996; Tennakoon et al., 2011; Lo Gullo et al., 2012) and *S. spicatum* had no effect on host foliar [Fe] (Struthers et al., 1986). Significant enrichment of [Fe] as a result of infection may lead to an excess of free radical production impairing cellular structure and damaging membranes, DNA and proteins (de Dorlodot et al., 2005).

## 5 CONCLUSION

We successfully established the native parasite, *C. pubescens*, on one of Australia’s worst weeds, *R. anglocandicans* at three field sites. We also demonstrated that *C. pubescens* significantly impacted *F*_v_/*F*_m_, [N] and [P]-status of *R. anglocandicans* across these sites. Mowing did not affect parasite impact on photosynthetic performance of *R. anglocandicans*, but did enhance negative parasite effects on host [N] at one site. The results support the potential application of *C. pubescens* as a native biocontrol on *R. anglocandicans* in congruence with this native parasite consistently negatively affecting other major invasive species and continue to suggest that native parasites can be effective weed biocontrols to help conserve and restore biodiversity. This work highlights that native parasitic plants should be incorporated into the theoretical frameworks of invasion theory, namely the Biotic Resistance theory for control of invasive species.

## ACKNOWLEDGEMENTS

Special thanks to Grace Porter-Dabrowski for assistance, and Bernardo O’Connor, Emily Reynolds and Hayley Jose for their support with fieldwork. Park Rangers at study sites and the DEW fire crew for mowing plots. This work was supported by the Department of Agriculture, Water and the Environment (Australian Government) [55119480] and Primary Industries and Regions SA (Biosecurity) [56119496].

## Author Contributions

*RMC and JMF conceived and designed the experiment. RMC performed the experiment and analysed the data. RMC, JMF, and JRW interpreted the analysis and wrote the manuscript.

