## Supplementary material for "Impact of a native hemiparasite and mowing on performance of a major invasive weed, European blackberry": Supp Tables S1-S4

### SUPPORTING INFORMATION

**TABLE S1** Unmown: two-way ANOVA results for the effects of infection with *Cassytha pubescens* (I) and site (S) on number of fruits, number of prickles, predawn and midday quantum yield ($F_v/F_m$ and $\Phi_{PSII}$), midday electron transport rate (ETR), foliar carbon isotope composition ($\delta^{13}C$), nitrogen [N], phosphorus [P], sodium [Na] and iron concentration [Fe] of *Rubus anglocandicans*

| | Fruit | Prickles | $F_v/F_m$ | $\Phi_{PSII}$ | ETR | $\delta^{13}C$ | [N] | [P] | [Na] | [Fe] |
|---|---|---|---|---|---|---|---|---|---|---|
| I | *9.45* | *8.91* | *19.2* | *36.7* | *35.4* | *0.044* | *24.8* | *18.3* | *50.7* | *45.8* |
| | 0.507 | 0.314 | 7.39 | 0.060 | 1.49 | 0.024 | 1.31 | *0.394* | *6.52* | *1.88* |
| S | *10.6* | *10.4* | *2.37* | *7.29* | *6.16* | *33.5* | *11.9* | *77.9* | *6.18* | *46.3* |
| | 1.14 | 0.731 | 1.82 | 0.024 | 0.519 | 37.3 | 1.26 | *3.36* | *1.59* | *3.80* |
| I × S | *3.34* | *0.702* | *0.327* | *11.3* | *12.0* | *6.22* | *0.218* | *2.27* | *1.73* | *8.86* |
| | 0.358 | 0.050 | 0.252 | 0.037 | 1.01 | 6.93 | 0.023 | *0.098* | *0.444* | *0.727* |
| Error | 2.47 | 1.62 | 20.8 | 0.089 | 2.27 | 30.1 | 2.86 | 1.16 | 6.05 | 2.22 |
| *df* | 1,2-46 | 1,2-46 | 1,2-54 | 1,2-54 | 1,2-54 | 1,2-54 | 1,2-54 | 1,2-54 | 1,2-47 | 1,2-54 |

*F* and sum of squares italic and regular type, respectively, and degrees of freedom (*df*). To meet model assumptions: fruit, $\Phi_{PSII}$ and ETR transformed to power of 0.25; prickle number, [P], [Na] and [Fe] log transformed; $F_v/F_m$ inverse log transformed.

**TABLE S2** Mowing effect: three-way ANOVA results for the effects of infection with *Cassytha pubescens* (I), site (S) and mowing (M) on predawn and midday quantum yield ($F_v/F_m$ and $\Phi_{PSII}$), midday electron transport rate (ETR), foliar carbon isotope composition ($\delta^{13}C$), nitrogen [N], phosphorus [P], sodium [Na] and iron concentration [Fe] of *Rubus anglocandicans*

| | $F_v/F_m$ | $\Phi_{PSII}$ | ETR | $\delta^{13}C$ | [N] | [P] | [Na] | [Fe] |
|---|---|---|---|---|---|---|---|---|
| I | *13.6* | *25.0* | *25.6* | *3.98* | *28.0* | *9.88* | *36.6* | *27.6* |
| | 5.37 | 0.071 | 46.7 | 2.02 | 1.42 | 0.247 | 4.58 | 1.65 |
| S | *11.7* | *21.4* | *26.3* | *125* | *29.3* | *259* | *52.5* | *120* |
| | 4.61 | 0.061 | 48.0 | 63.6 | 1.49 | 6.48 | 6.57 | 7.19 |
| I × S | *0.359* | *13.3* | *16.6* | *5.26* | *1.40* | *1.95* | *4.22* | *13.8* |
| | 0.142 | 0.038 | 30.3 | 2.67 | 0.071 | 0.049 | 0.528 | 0.829 |
| M | *1.49* | *0.040* | *0.140* | *19.5* | *32.0* | *13.4* | *10.8* | *14.1* |
| | 0.586 | 0.0001 | 0.255 | 9.89 | 1.63 | 0.334 | 1.35 | 0.842 |
| I × M | *1.28* | *3.17* | *2.34* | *0.002* | *0.843* | *0.071* | *0.390* | *1.82* |
| | 0.505 | 0.009 | 4.27 | 0.001 | 0.043 | 0.002 | 0.049 | 0.109 |
| S × M | *1.64* | *2.99* | *6.69* | *2.43* | *1.28* | *1.97* | *12.8* | *0.808* |
| | 0.646 | 0.009 | 12.2 | 1.23 | 0.065 | 0.049 | 1.60 | 0.049 |
| I × S × M | *0.278* | *2.32* | *1.79* | *1.43* | *4.18* | *0.161* | *0.288* | *0.006* |
| | 0.110 | 0.007 | 3.28 | 0.728 | 0.213 | 0.004 | 0.036 | 0.0003 |
| Error | 25.2 | 0.176 | 113 | 31.5 | 3.15 | 1.55 | 6.51 | 3.72 |
| *df* | 1-64 | 1-62 | 1-62 | 1-62 | 1-62 | 1-62 | 1-52 | 1-62 |

*F* and sum of squares italic and regular type, respectively, and degrees of freedom (*df*). To meet model assumptions: $F_v/F_m$ inverse log transformed; $\Phi_{PSII}$, and ETR square root transformed; [P], [Na] and [Fe] log transformed.

**TABLE S3** ANOVA results for effects of site (S) on parasite predawn and midday quantum yield ($F_v/F_m$ and $\Phi_{PSII}$) and midday electron transport rates (ETR) of *Cassytha pubescens* infecting *Rubus anglocandicans*. Similarly for mown (M) and unmown sites

| | $F_v/F_m$ | $\Phi_{PSII}$ | ETR |
|---|---|---|---|
| *Unmown* | | | |
| S | **0.020*** | **0.003**** | **0.0008***** |
| | *4.57* | *7.54* | *9.40* |
| | 3.52 | 1.09 | 1.40 |
| Error | 10.4 | 1.96 | 2.01 |
| *Mowing effect* | | | |
| S | **0.675** | **<0.0001***** | **<0.0001***** |
| | *0.179* | *47.4* | *58.1* |
| | 0.056 | 4.00 | 5.35 |
| M | **0.118** | **0.487** | **0.187** |
| | *2.59* | *0.495* | *1.83* |
| | 0.803 | 0.042 | 0.168 |
| S × M | **0.207** | **0.002**** | **0.0006** |
| | *1.66* | *11.2* | *14.8* |
| | 0.515 | 0.947 | 1.36 |
| Error | 9.32 | 2.53 | 2.76 |

***p***, *F* and sum of square values are in bold, italic and regular type, respectively. Degrees of freedom: Unmown = 2, 27 for all variables; Mowing effect = 1, 30 for all variables. Significance codes: 0'***', 0.001'**', 0.01'*'. To meet model assumptions: $F_v/F_m$ inverse log transformed and $\Phi_{PSII}$ and ETR log transformed.

**TABLE S4** ANOVA results for the effects of infection with *Cassytha pubescens* (I) on stomatal conductance ($g_s$) of *Rubus anglocandicans*. Similarly for mown (M) and unmown sites

| $g_s$ | Belair | Horsnell | Blackwood |
|---|---|---|---|
| *Unmown* | | | |
| I | *22.0* | *2.15* | *1.26* |
| | 0.0003 | 3465 | 7767 |
| Error | 0.0001 | 16106 | 61592 |
| *Mowing effect* | | | |
| I | N/A | *10.3* | *1.51* |
| | | 22016 | 0.328 |
| M | N/A | *9.57* | *13.3* |
| | | 20446 | 2.88 |
| I × M | N/A | *1.99* | *0.175* |
| | | 4243 | 0.038 |
| Error | N/A | 42713 | 4.34 |

*F* and sum of square values are in italic and regular type, respectively. Degrees of freedom: Unmown = 1, 10 for all sites and Belair data inverse transformed; Mowing effect = 1, 20 for both sites and Blackwood data transformed to the power of 0.25. N/A = not applicable.
