## Supplementary material for "Impact of a native hemiparasite and mowing on performance of a major invasive weed, European blackberry": Supp Figs S1-S9

**SUPPORTING INFORMATION**

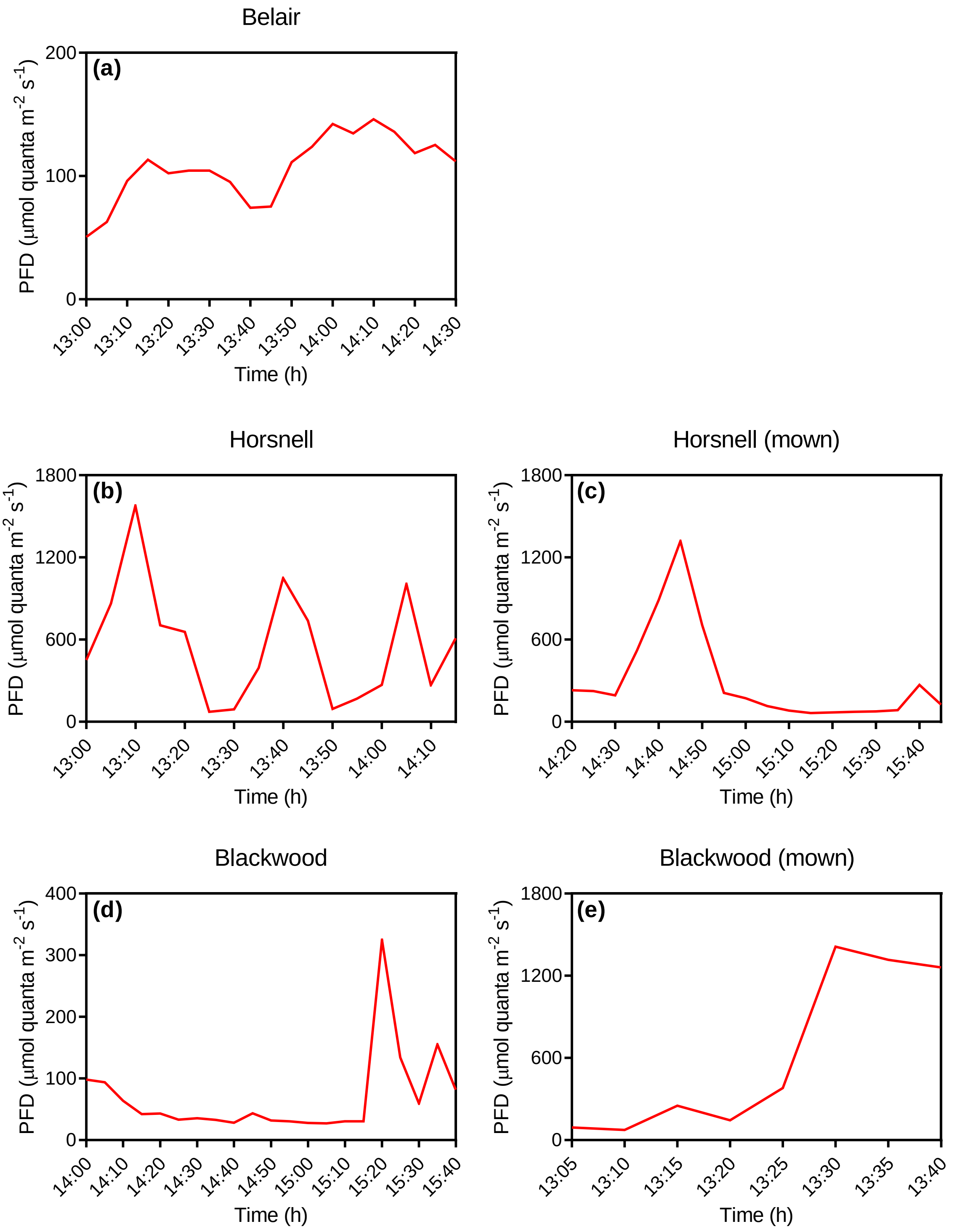


**FIGURE S1** Photo flux density during midday quantum yield ($\Phi_{PSII}$) and midday electron transport measurements on unmown or mown *Rubus anglocandicans* and *Cassytha pubescens* at Belair (27/03/2019), Horsnell (02/04/2019) and Blackwood (28/03/2019) in the Mt Lofty Ranges of South Australia

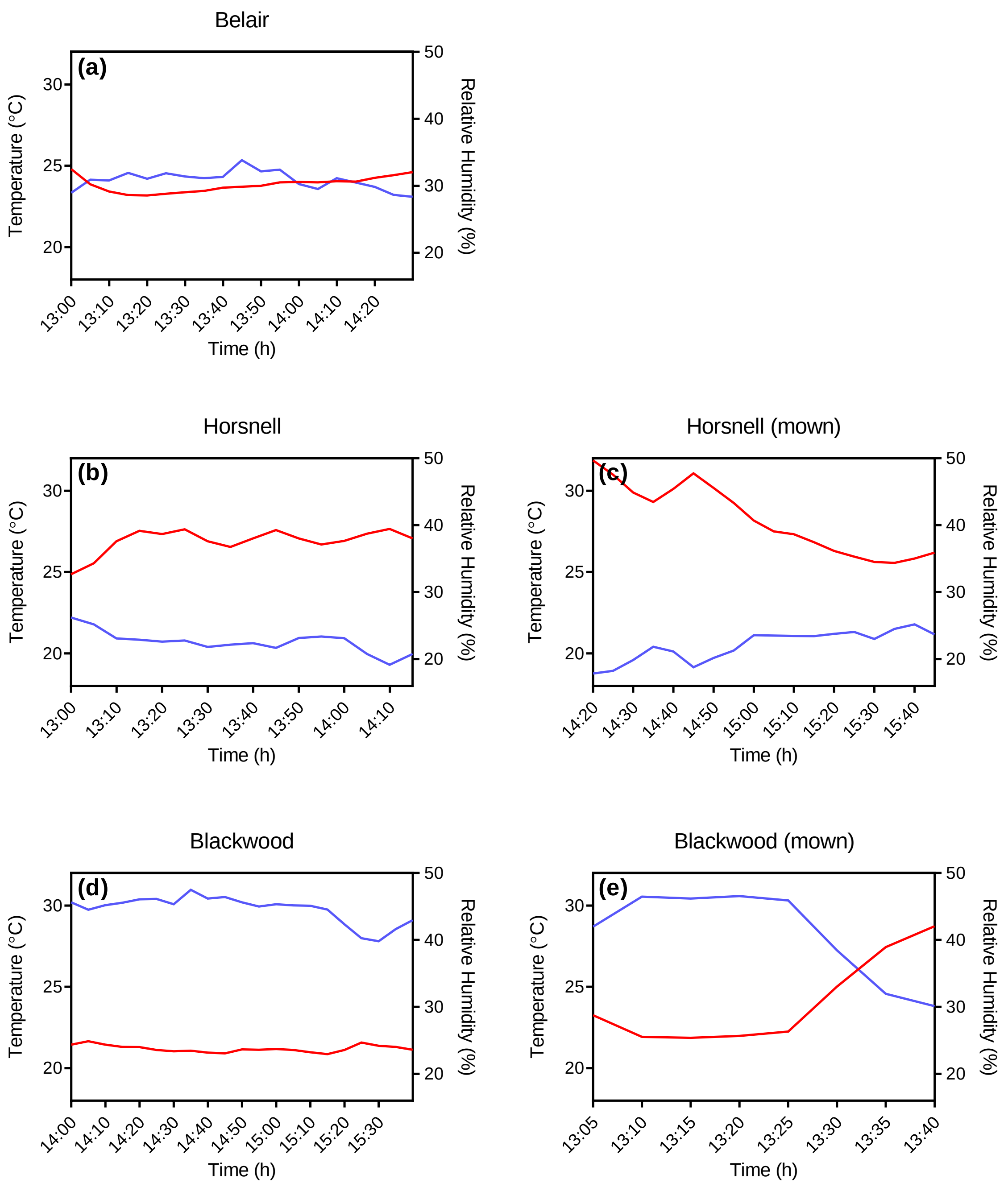


**FIGURE S2** Air temperature (left-axis: red line) and relative humidity (right-axis: blue line) during midday quantum yield ($\Phi_{PSII}$) and midday electron transport measurements on unmown or mown *Rubus anglocandicans* and *Cassytha pubescens* at Belair (27/03/2019), Horsnell (02/04/2019) and Blackwood (28/03/2019) in the Mt Lofty Ranges of South Australia

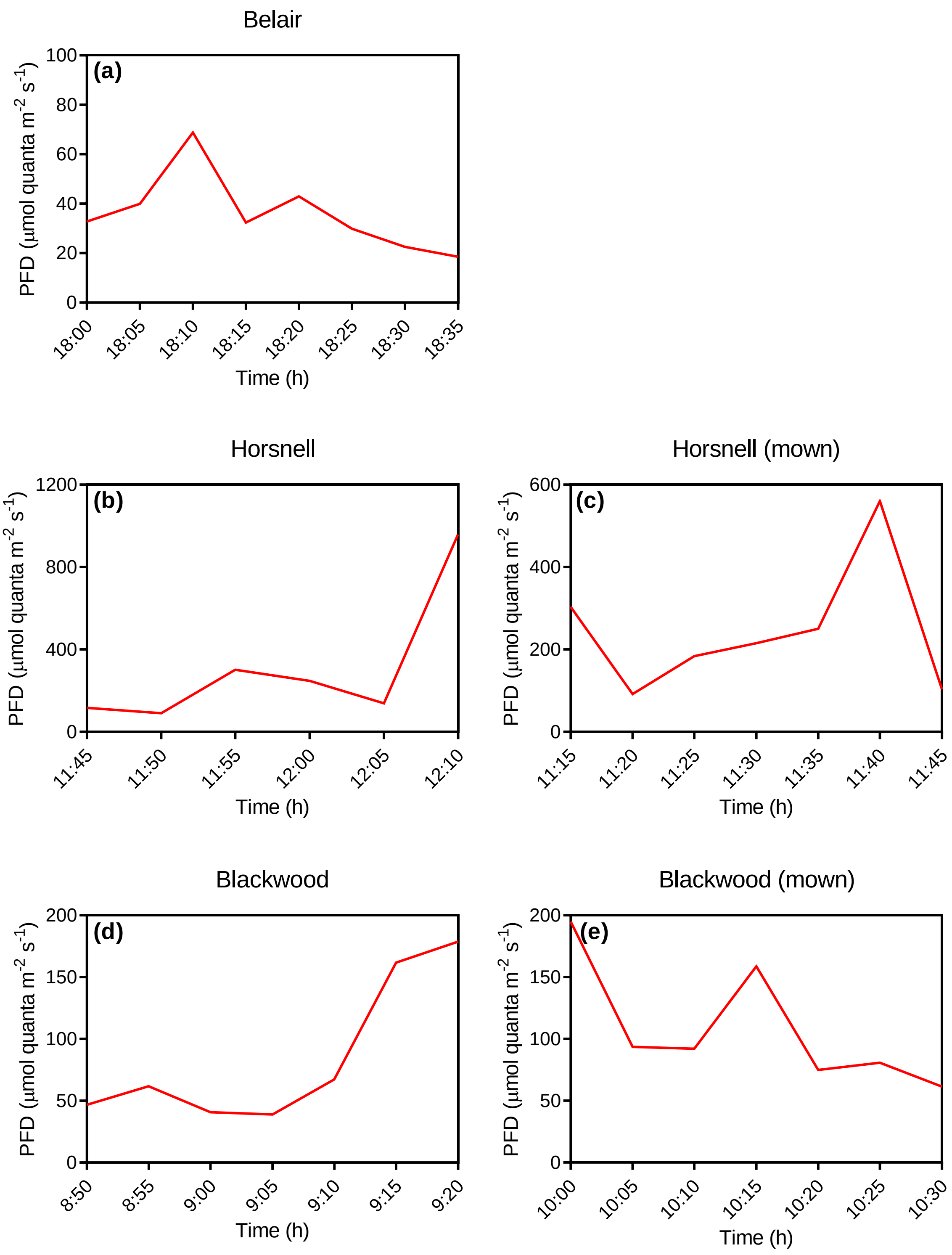


**FIGURE S3** Photo flux density during porometer measurements on unmown or mown *Rubus anglocandicans* and *Cassytha pubescens* at Belair (04/04/2019), Horsnell (11/04/2019) and Blackwood (07/04/2019) in the Mt Lofty Ranges of South Australia

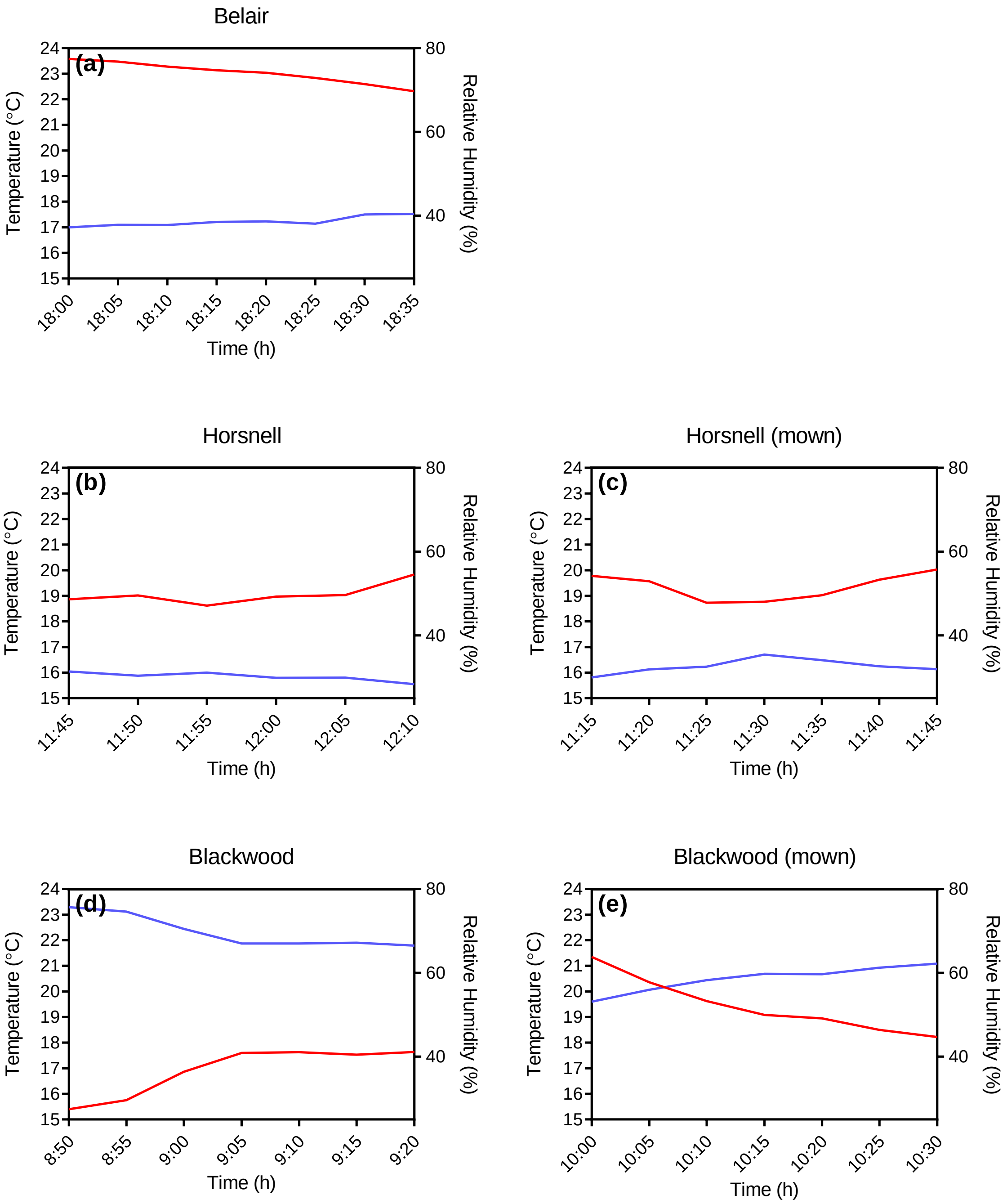


**FIGURE S4** Air temperature (left-axis: red line) and relative humidity (right-axis: blue line) during midday quantum yield ($\Phi_{PSII}$) and midday electron transport measurements on unmown or mown *Rubus anglocandicans* and *Cassytha pubescens* at Belair (04/04/2019), Horsnell (11/04/2019) and Blackwood (07/04/2019) in the Mt Lofty Ranges of South Australia

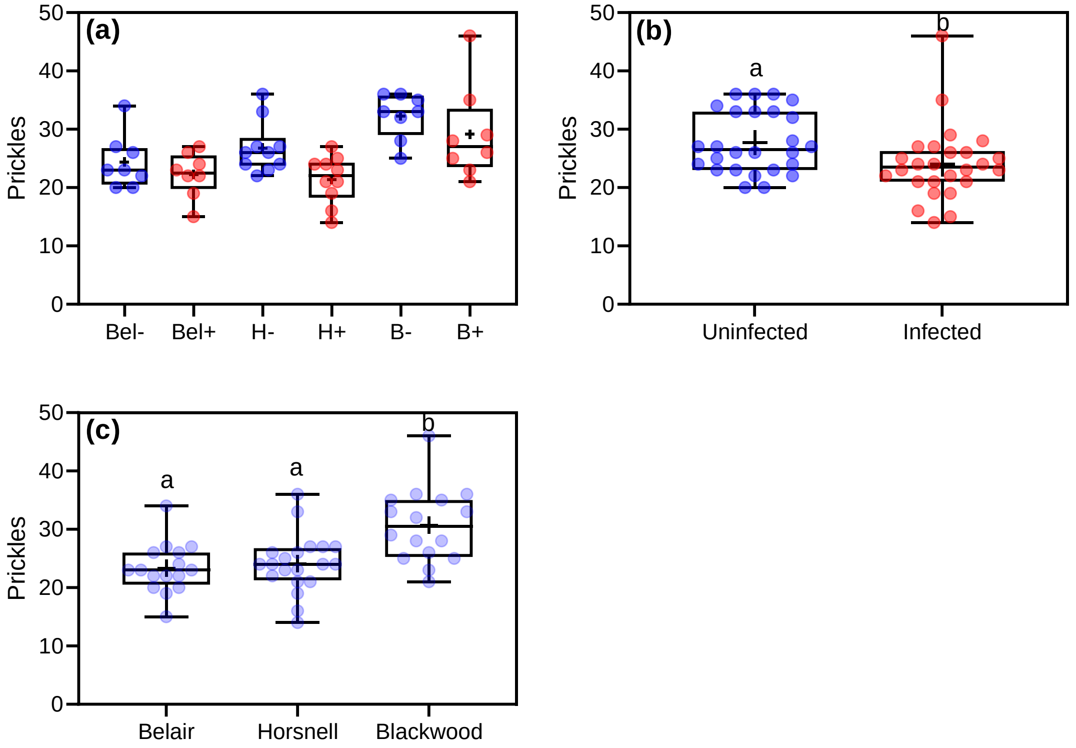


**FIGURE S5** (a) Number of prickles per cane (30 cm from cane tip) of *Rubus anglocandicans*, when uninfected (–) or infected (+) with *Cassytha pubescens* at Belair (Bel–, Bel+), Horsnell (H–, H+) and Blackwood (B–, B+), respectively. Main effects of (b) infection and (c) site on host prickle number per cane. All data points, median, percentile lines and mean (+ within box) are displayed, different letters indicate significant differences: (a) $n$ = 8–10, (b) $n$ = 26, and (c) $n$ = 16–20

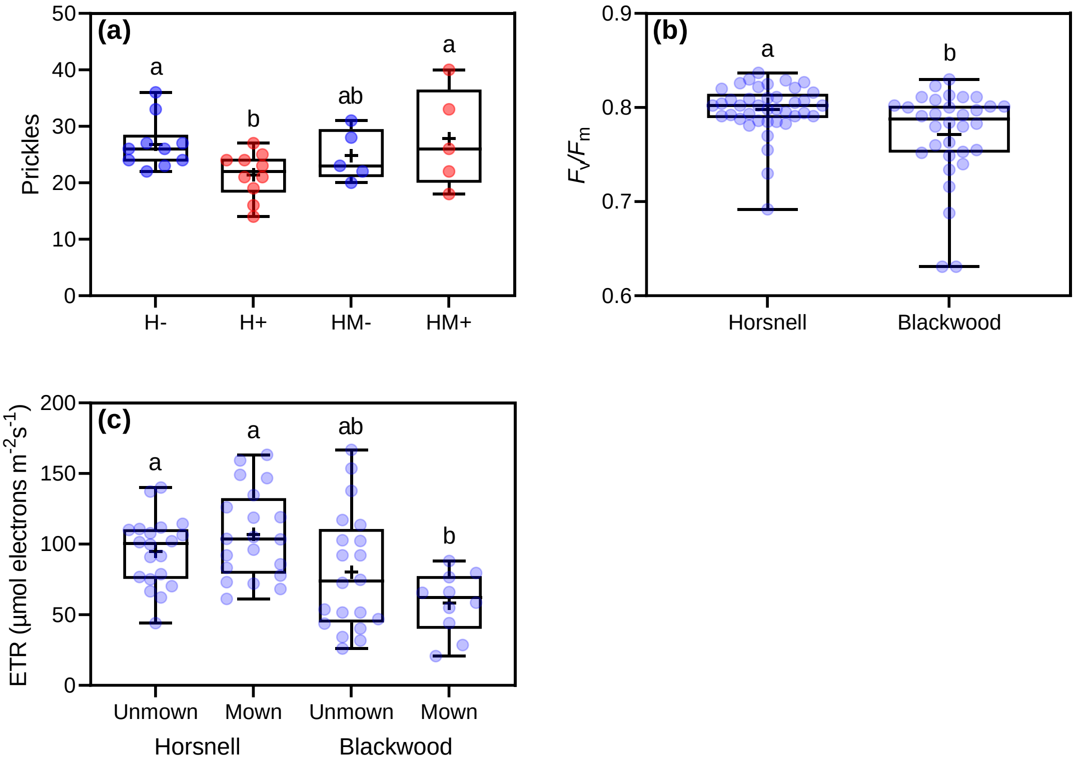


**FIGURE S6** (a) Number of prickles per cane (30 cm from cane tip) of *Rubus anglocandicans*, when unmown or mown (m) and uninfected (–) or infected (+) with *Cassytha pubescens* at Horsnell (unmown: H–, H+, mown: HM–, HM+) and Blackwood (unmown: B–, B+, mown: BM–, BM+), respectively. (b) Main effect of infection on host predawn quantum yield ($F_v/F_m$). (c) Site × mowing interaction on host midday electron transport rates (ETR). All data points, median, percentile lines and mean (+ within box) are displayed, different letters indicate significant differences: (a) *n* = 5–10, (b) *n* = 32–40, (c) *n* = 10–20 and (d) *n* = 40 and 30 (unmown and mown, respectively)

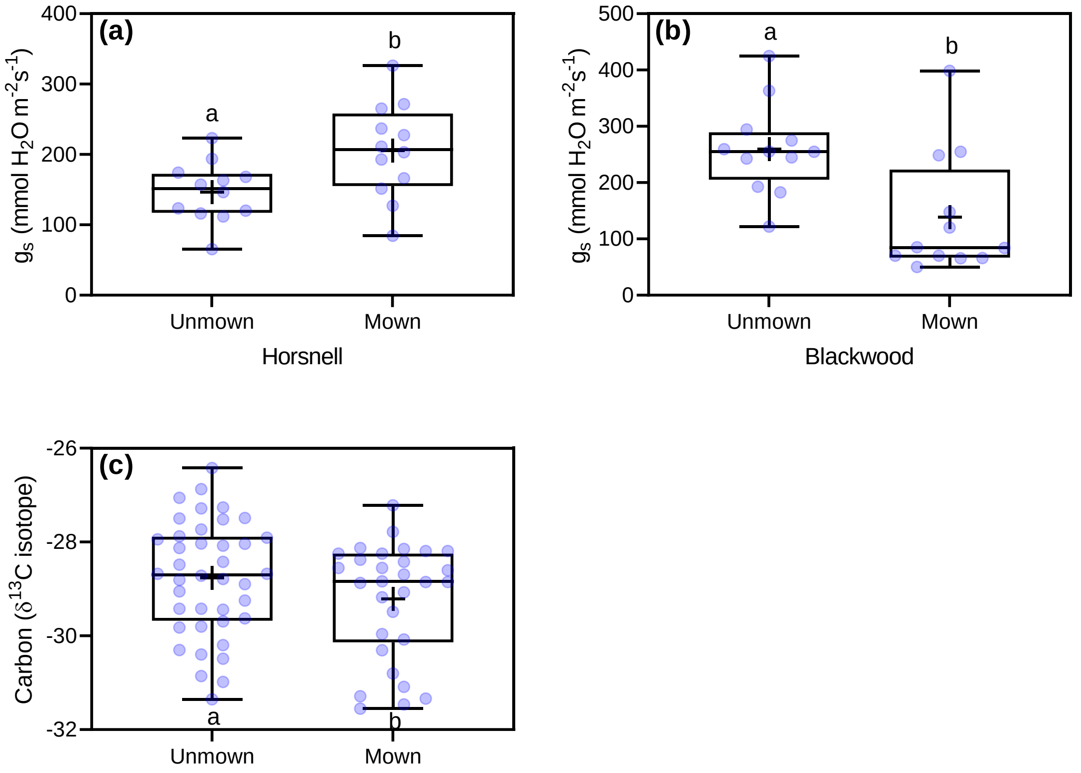


**FIGURE S7** Main effect of mowing on stomatal conductance ($g_s$) of *Rubus anglocandicans* at (a) Horsnell and (b) Blackwood. (c) Main effect of mowing on carbon isotope composition of the host. All data points, median, percentile lines and mean (+ within box) are displayed, different letters indicate significant differences: (a, b) $n$ = 12 and (c) $n$ = 40 and 30 (unmown and mown, respectively)

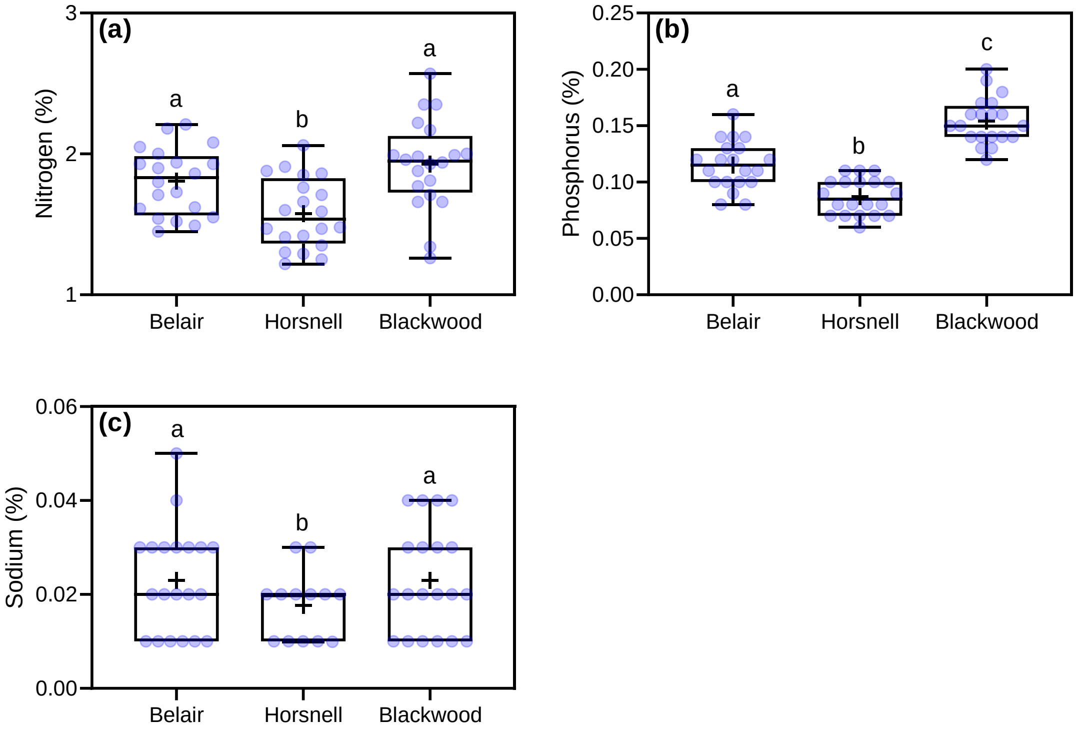


**FIGURE S8** Main effect of site on leaf (a) nitrogen, (b) phosphorus and (c) sodium concentration of *Rubus anglocandicans* at Belair, Horsnell and Blackwood. All data points, median, percentile lines and mean (+ within box) are displayed, different letters indicate significant differences: (a, b) $n$ = 20 and (c) $n$ = 13

.

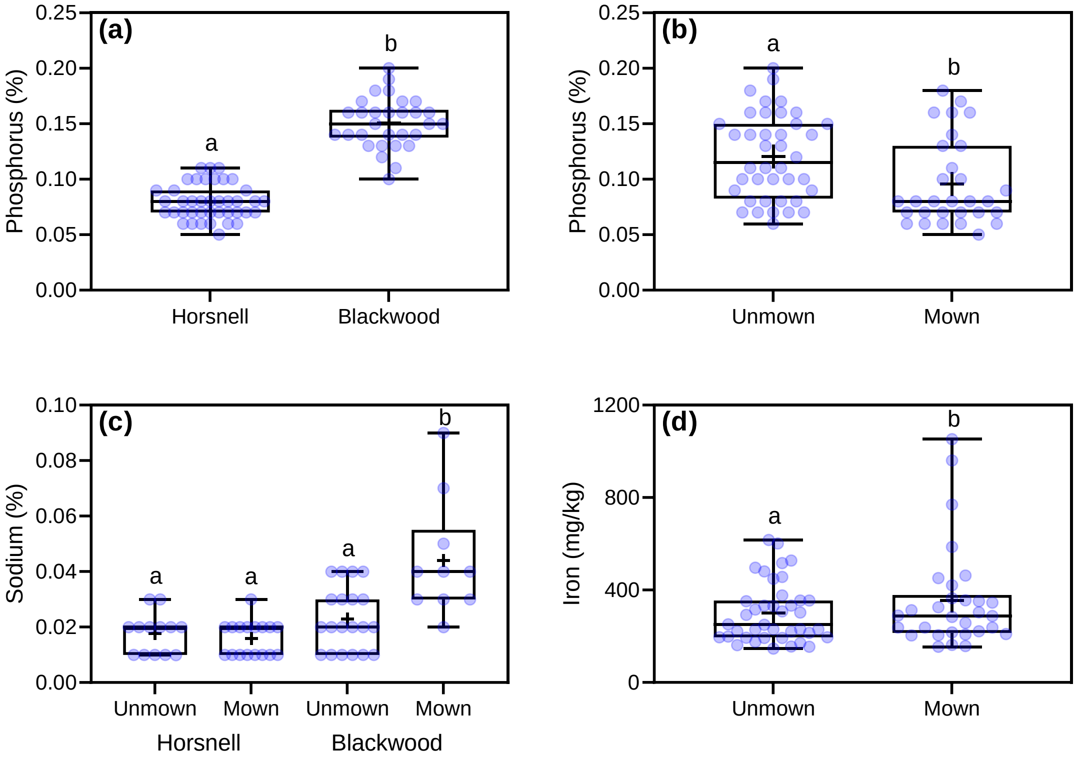


**FIGURE S9** (a) Main effect of site on leaf nitrogen, and main effect of mowing on leaf (b) phosphorus and (d) sodium concentration of *Rubus anglocandicans* at Horsnell and Blackwood. (c) Site × mowing effect on host sodium concentration. All data points, median, percentile lines and mean (+ within box) are displayed, different letters indicate significant differences: (a) $n$ = 40 and 30 (Horsnell and Blackwood, respectively), (b, d) $n$ = 40 and 30 (unmown and mown, respectively) and (c) $n$ = 10–20
